# EEG dynamics in Lewy body diseases is related to clinical fluctuations and degeneration of locus coeruleus

**DOI:** 10.64898/2026.01.20.700527

**Authors:** Eva Výtvarová, Kristína Mitterová, Marianna Angiolelli, Anežka Kovářová, Alžběta Šejnoha Minsterová, Luboš Brabenec, Martin Lamoš, Pierpaolo Sorrentino, Irena Rektorová, Jan Fousek

## Abstract

This work investigates aperiodic bursts of neuronal activity, referred to as neuronal avalanches, which are thought to support flexible information processing in Lewy body (LB) disease. We compared avalanche-derived features across LB disease phenotypes (prodromal dementia with LB and Parkinson’s disease, PD), with healthy controls (HC). Based on the premise that focal neurodegeneration alters whole-brain dynamics, we examined whether the “richness” of braindynamics was associated with clinical symptoms and with the microstructural integrity of catecholaminergic nuclei - the right caudal locus coeruleus (rcLC) and the substantia nigra pars compacta (SNpc).

The sample comprised 30 HC, 56 cognitively normal individuals with core clinical features of dementia with LB (CN-CCF), 34 individuals with mild cognitive impairment with LB (MCI-LB), and 30 advanced PD patients. Neuronal avalanches were extracted from source-reconstructed resting-state EEG and described by their count, the number of unique spatial patterns (flexibility), hemispheric symmetry of flexibility, and avalanche transition matrices. LC and SNpc integrity were assessed using neuromelanin-sensitive MRI and free-water diffusion imaging, respectively.

Lewy body disease groups exhibited increased flexibility relative to HC, which was, in prodromal groups, associated with cognitive fluctuations and REM sleep behavior disorder symptoms, but not parkinsonism. Flexibility was negatively related to rcLC integrity, which was lowest in MCI-LB. These findings suggest that elevated flexibility reflects dysregulated brain dynamics linked to malignant non-motor features of LB diseases and to the caudal LC involvement.

## 1. Introduction

Dementia with Lewy bodies (DLB) is a progressive neurological disorder characterized by a multidomain cognitive decline [1, 2] accompanied by noncognitive symptoms— notably isolated REM sleep behavioral disorder, parkinsonism, visual hallucinations, and fluctuating cognition and alertness [1, 3, 4]. Together with Parkinson’s disease (PD) it belongs to the group of Lewy body diseases (LBD), which are characterized by early synaptic disruption impacting functional and structural connectivity [5]. On the spectrum of LBDs, early PD is a less malignant disease subtype predominantly characterized by asymmetric motor symptoms of parkinsonism while DLB is a more malignant subtype mostly characterized by fluctuating cognition progressing to early dementia [6].

In LBD, *α*-synuclein–related synaptic dysfunction is regionally localized yet accompanied by widespread large-scale alterations, making it essential to examine intrinsic neural dynamics, defined as spontaneous activity emerging from ongoing neuronal interactions, to understand how focal pathology propagates through distributed circuits and leads to cognitive fluctuations and other clinical features of LBD. During rest and task performance, multiple brain regions are recruited within the brain networks in simultaneous activation sequences that underpin specific functions [7]. Brain activity is characterized by bursts of activation, in which neural signals intermittently exceed a certain threshold [8].

These transient periods of increased activity are termed cascades, or neuronal avalanches, when their size and duration distributions follow a power-law, a defining hallmark of critical brain dynamics [9]. Each neuronal avalanche is characterized by a specific spatiotemporal activation pattern, defined by the set of brain regions recruited during the event. The total number of distinct avalanche patterns observed over time is referred to as *flexibility* [10].

In the healthy brain, high flexibility reflects the dynamic nature of neural activity, enabling a rich variety of avalanche patterns that recruit different regions and give rise to a rich functional repertoire and less stereotyped activity. This diversity supports efficient and adaptive information processing across whole-brain network. In contrast, in neurodegenerative conditions, the flexibility of the brain can be either decreased - reflecting a reduced accessible functional repertoire - or increased, pointing to presence of pathological activity patterns or a general disregulation of spontaneous neural fluctuations dynamics. Reduced flexibility has been associated with PD [11], amyotrophic lateral sclerosis [10], or amnestic mild cognitive impairment (MCI) [12]. On the other hand, increased flexibility was observed in relapsing–remitting but not in secondary progressive multiple sclerosis [13].

While flexibility captures the diversity of activation patterns, it does not account for the temporal organization of these patterns. Brain activity is inherently sequential, and the functional relevance of neuronal avalanches lies not only in the variety of patterns that emerge, but also in how the system transitions between them. Consequently, a purely static description of the avalanche repertoire can be improved by investigating the transition probabilities between neuronal avalanche patterns. These probabilities provide a dynamical description of the exploration of the state space [14]. They and derived characteristics have been shown to be influenced in epilepsy [15], amnestic MCI [12], by levodopa in PD [16], or by task [7], and they have been used as an informative data feature in building the virtual Parkinsonian patient model [17]. The analysis of avalanche transition probabilities thus complements flexibility by revealing the constraints and rules governing the temporal evolution of neuronal activity. Together, these measures offer a more complete characterization of brain dynamics, bridging the gap between the richness of the activation repertoire and the structure.

The present work focused on examining measures derived from neuronal avalanches in prodromal dementia with Lewy bodies (DLB), in the context of prior findings indicating altered functional connectivity and brain dynamics [18]. Specifically, we assessed whether avalanche patterns in prodromal DLB differ from those in HC and advanced PD, and whether such differences reflect previously observed compensatory processes [19] or rather are related to fluctuations of alertness.

As multiple neurotransmitter systems are affected in Lewy body diseases (LBD), we examined the relationship between the integrity of locus coeruleus (LC) and substantia nigra pars compacta (SNpc), and unique avalanche patterns (that is, flexibility). These two catecholamine systems are one of the earliest affected in prodromal LBD. In particular, early pathology in LC pathology, one of the major modulators of brain activity, is thought to precede degeneration in the SNpc, particularly in patients exhibiting REM sleep behavior disorder, visual hallucinations, or fluctuations of cognition. By contrast, the SNpc, a dopaminergic nucleus with relatively more concentrated projections, primarily supports motor, associative, and reward-related circuits [6]. A simultaneous, multimodal MRI assessment of the microstructural integrity of these catecholaminergic nuclei may therefore provide deeper insight into the origins of network alterations in early LBD subtypes.

We hypothesized that (H1) HC will differ from LBD groups (cognitively unimpaired, CN-CCF; mildly cognitively impaired, MCI-LB; and advanced PD) in flexibility. Converging evidence indicates that structural brain integrity constrains and shapes functional brain dynamics, such that preserved anatomical connectivity supports flexible patterns of neural activity. In healthy conditions, a positive correlation between flexibility—derived from neuronal avalanche dynamics—and brain connectivity has been previously demonstrated [14]. Against this background, and based on empirical evidence linking *α*-synuclein–related symptomatology to disruptions of the brain connectome [20], we hypothesized that (H2) in prodromal DLB and advanced PD, flexibility will be associated with clinical features of LBD, such as fluctuating cognition, symptoms of REM sleep behavioral disorder, and parkinsonism [1, 6] and to structural integrity of the LC (particularly its right caudal subregion) [21] and SNpc.

Lastly, because early motor symptoms in PD are typically lateralized to one side of the body [22], we introduced a novel measure of hemispheric asymmetry based on neuronal avalanche flexibility. Specifically, hemispheric asymmetry is quantified as the ratio between right- and left-hemisphere flexibility, scaled between 0 and 1, where 0 indicates no asymmetry and 1 indicates maximal asymmetry. We hypothesized (H3) that greater hemispheric asymmetry in flexibility will be associated with motor impairment in the prodromal DLB and PD groups. Finally, we predicted that (H4) HC and LBD groups will differ in avalanche transition probabilities [17].

## 2 Results

### 2.1 Sample and descriptive statistics

Our sample of 150 subjects comprised 56 CN-CCF, 34 MCI-LB, 30 HC, and 30 advanced PD. The prodromal DLB groups (CN-CCF, MCI-LB) did not significantly differ from HC in terms of age and sex, but had significantly lower education and integrity of the rcLC, higher flexibility (= the number of unique avalanche patterns observed over time), core clinical features of DLB, and depressive symptoms (Table 1). On the other hand, the integrity of SNpc did not significantly differ across groups.

**Table 1.** Sample description. HC – cognitively healthy; CN-CCF – cognitively unimpaired, and MCI-LB – mildly cognitively impaired subjects with prodromal LBD; PD – patients with Parkinson’s disease; Q1, Q3 – 25^th^ and 75^th^ percentile; *p* – *p*-values from the Mann-Whitney *U* test or the *χ*^2^ – Chi-square test; f/m – females vs. males; MoCA – Montreal cognitive assessment [23]; rcLC* – right caudal locus coeruleus; lSNpc/rSNpc* - left/right substantia nigra pars compacta; UPDRS – Movement Disorders Society - Unified Parkinson’s Disease Rating Scale part 3 [24]; MFS – Mayo Fluctuations Scale [25]; RBDq – REM Sleep Behavior Disorder questionnaire [26]; NPI – Neuropsychiatric Inventory [27]; GDS* - Geriatric Depression Scale [28]; LEDD – levodopa equivalent daily dose; *scales and MRI were not available for subjects with PD.

| | Median (Q1, Q3) or count | | | | $p$ (U or $\chi^2$) HC versus groups | | |
| --- | --- | --- | --- | --- | --- | --- | --- |
|  | HC<br>(n=30) | CN-CCF<br>(n=56) | MCI-LB<br>(n=34) | PD<br>(n=30) | HC vs.<br>CN-CCF | HC vs.<br>MCI-LB | HC vs.<br>PD |
| Age | 68 (62, 74) | 66 (61, 70) | 70 (65, 73) | 68 (61, 76) | .12 | .58 | .93 |
| Sex (f/m) | 16/14 | 34/22 | 22/13 | 9/21 | .65 | .46 | .07 |
| Education | 17 (14, 19) | 15 (12, 18) | 13 (12, 15) | 18 (15, 18) | .06 | < .001 | .68 |
| MoCA | 27 (26, 29) | 27 (26, 28) | 24 (22, 26) | 26 (24, 27) | .50 | < .001 | .002 |
| Flexibility | 526 (376, 681) | 743 (549, 936) | 766 (542, 1028) | 689 (521, 875) | .002 | .011 | .017 |
| Flexibility<br>symmetry | .11(.02, .18) | .13 (.08, .23) | .11 (.05, .18) | .12 (.08, .20) | .07 | .60 | .33 |
| rcLC* | .14 (.10, .18) | .11 (.08, .17) | .09 (.07, .14) | - | .12 | .003 | - |
| lSNpc* | .29 (.26, .31) | .29 (.26, .33) | .26 (.20, .29) | - | .90 | .20 | - |
| rSNpc* | .29 (.26, .31) | .30 (.27, .35) | .28 (.26, .31) | - | .21 | .78 | - |
| UPDRS | 0 (0, 1.5) | 4 (2.5, 6.5) | 5 (3, 8) | 11 (8, 15) | < .001 | < .001 | < .001 |
| MFS | 0 (0, 0) | 1 (0, 1) | 1 (0, 2) | 0 (1, 2) | < .001 | < .001 | < .001 |
| RBDq | 1 (0, 3) | 4 (2, 6) | 5 (2.5, 5.5) | 4 (3, 7) | < .001 | < .001 | < .001 |
| NPI (+/-) | 0/33 | 10/46 | 6/30 | 4/19 | .01 | .03 | .024 |
| GDS* | 3 (1, 6) | 7.5 (4, 11.5) | 8 (5, 11) | - | < .001 | < .001 | - |
| LEDD | - | - | - | 840 (703, 1320) | - | - | - |

PD had significantly lower cognitive impairment (*p* = .021), but more severe parkinsonism (the UPDRS score, *p* < .001) than MCI-LB; and higher cognitive impairment (the MOCA score, *p* = .007) and higher UPDRS score (*p* = .001) than CN-CCF and HC. Other core clinical features, such as the fluctuations in cognition (MFS), symptoms of REM sleep behavior disorder (RBDq), and visual hallucinations (NPI) were not significantly different between advanced PD and both prodromal DLB groups. This suggests that while PD patients also manifested fluctuations in cognition and symptoms of RBD because of their progression, they had the highest degree of motor dysfunction.

### 2.2 Group differences in flexibility

A PERMANOVA [29] (20,000 iterations, Mahalanobis distance) was conducted on flexibility, with age, sex, and LEDD (in the case of PD) regressed out beforehand. The analysis revealed that flexibility (i.e., the number of unique avalanche patterns) differed between groups (*F* = 5.15, CI [.07, 3.23], *p* = .002; see Fig 1) and was significantly higher (*p* < .05, FDR corrected) in CN-CCF, MCI-LB (*F* = 11.81, CI [0, 5.36], *p* = .001; *F* = 12.46, CI [0, 5.40], *p* = .001, respectively), and uncorrected in the PD group (*F* = 6.65, CI [0, 5.30], *p* = .013) as compared to HC. Neither groups significantly differed in avalanche counts (i.e., the total number of avalanches occurring during the timeframe; *F* = 1.62, CI [.07, 3.29], *p* = .192; Supplementary Fig 4).

**Fig. 1.**
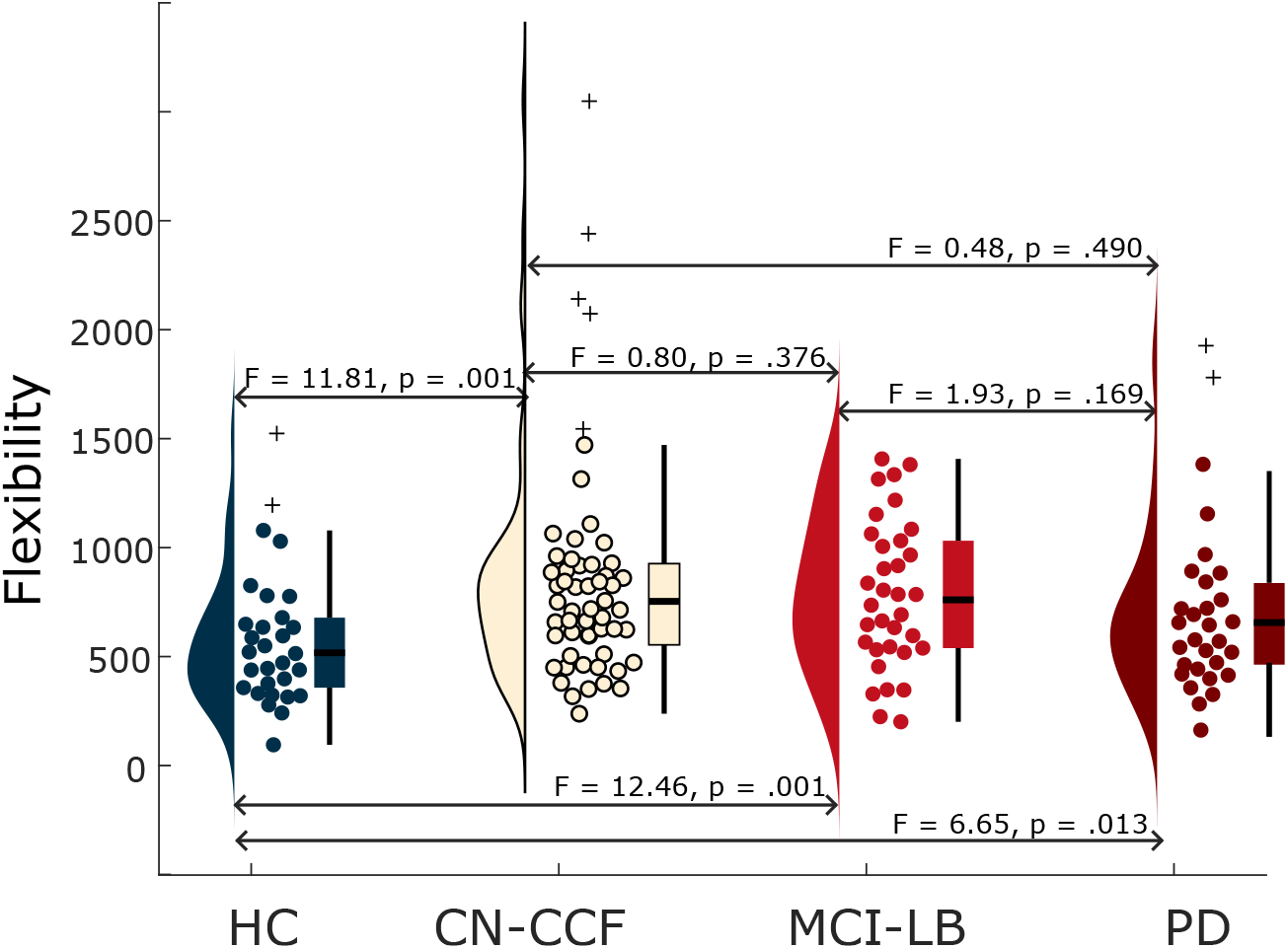
Group differences in flexibility. Note: Crosses (+) mark 9 outliers that were removed before statistical analysis; n(HC) = 28, n(CN-CCF) = 51, n(MCI-LB) = 34, n(PD) = 28). The half-violin plots, scatterplots, and boxplots were constructed using the daviolinplot function in Matlab [30]. Results for the analysis including outliers are reported in Supplementary Table 4.

### 2.3 Relationship between flexibility and clinical symptoms

Spearman’s partial correlations (age and sex as covariates, LEDD (in the case of PD) regressed out beforehand) demonstrated that greater flexibility was significantly associated with higher cognitive fluctuations (MFS) and greater symptoms severity of REM sleep behavior disorder (RBDq) across the full cohort (*p* < .05, FDR corrected; Table 2) and specifically within the MCI-LB group. Motor impairment was not related to flexibility. Further partial correlations controlling for Geriatric Depression Scale supported that these associations are invariant on the level of depressive symptoms (see Supplementary Table 5).

**Table 2.** Spearman’s partial correlations between flexibility and clinical features of interest. * *p* < .05, FDR corrected.

|  | UPDRS | MFS | RBDq |
| --- | --- | --- | --- |
| All subjects | $\rho = .13, p = .129$ | $\rho = .21, p = .012^*$ | $\rho = .22, p = .009^*$ |
| CN-CCF | $\rho = -.07, p = .596$ | $\rho = .09, p = .536$ | $\rho = -.03, p = .846$ |
| MCI-LB | $\rho = -.16, p = .395$ | $\rho = .38, p = .034$ | $\rho = .37, p = .035$ |
| PD | $\rho = -.11, p = .727$ | $\rho = -.19, p = .403$ | $\rho = -.08, p = .637$ |

### 2.4 Flexibility’s hemispheric symmetry

There was no significant difference in hemispheric symmetry of flexibility (*flexi ratio*) across all groups (see descriptive Table 1). However, Spearman’s partial correlations (age, sex, and LEDD (in the case of PD) as covariates) with the clinical features of interest (MFS and RBDq as those significantly related to the whole-brain flexibility), revealed a negative relationship between *flexi ratio* and MFS in MCI-LB group (n = 34; *rho* = −.36, *p* = .043), indicating that greater hemispheric symmetry in flexibility is associated with more pronounced cognitive fluctuations. Motor impairment was not related to hemispheric symmetry in either group (in PD: *rho* = −.35, *p* = .118).

### 2.5 Associations of neuromodulatory nuclei with flexibility

Spearman’s correlations between flexibility and rcLC (age and sex regressed out beforehand from both) yielded a weak, but significant negative relationship across HC and prodromal DLB groups (n = 100; excluding PD who did not have LC imaging), indicating that flexibility is inversely related to rcLC integrity (rho = − .21, *p* = .036; see Fig 2). The significance of this result was supported by a bootstrap correlation testing of 1,000 iterations on 50 randomly selected subjects; see Supplementary Fig 6).

**Fig. 2.**
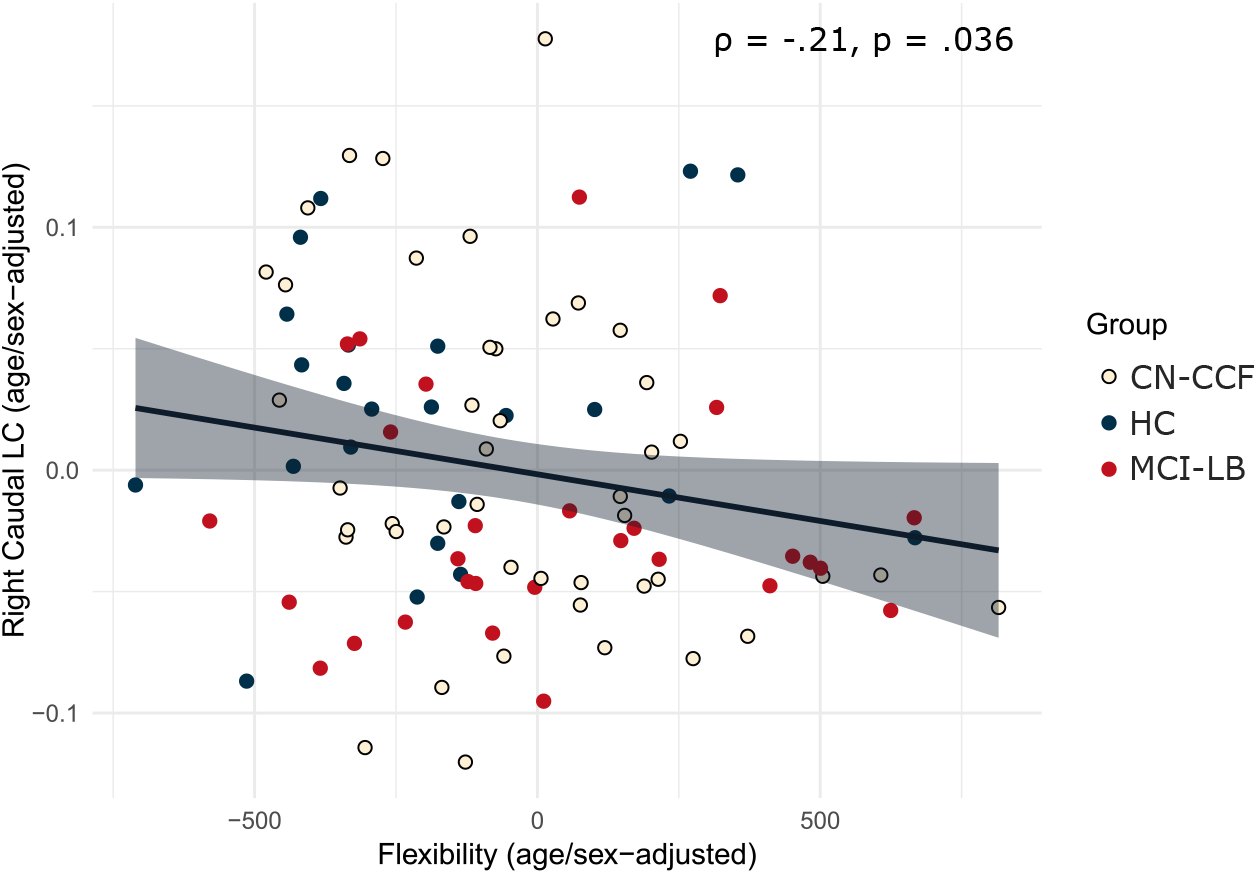
Relationship between flexibility and the right caudal locus coeruleus. Spearman’s correlation, age and sex regressed out. Plot was created in R using `ggplot2` [31].

Consistently with these results, Table 1 and Supplementary Fig 5 show significant groups differences in the rcLC integrity (*F* = 4.26, CI [.03, 3.85], *p* = .017), with significantly lower rcLC neuromelanine-signal intensity in the MCI-LB compared to HC.

On the other hand, flexibility did not correlate with volume of extracellular water of the left (rho = .03, *p* = .78) or right (rho = .04, *p* = .75) SNpc, indicating that flexibility and SNpc microstructural integrity are not related.

### 2.6 Avalanche patterns

To illustrate spatial patterns of avalanches, group-level avalanche transition matrices (ATMs) were obtained. By concatenating all avalanches from subjects within each group, the average transition probabilities across all these avalanches were computed and visualized in a matrix form and in a brain space [14]; see Fig 3a.

**Fig. 3.**
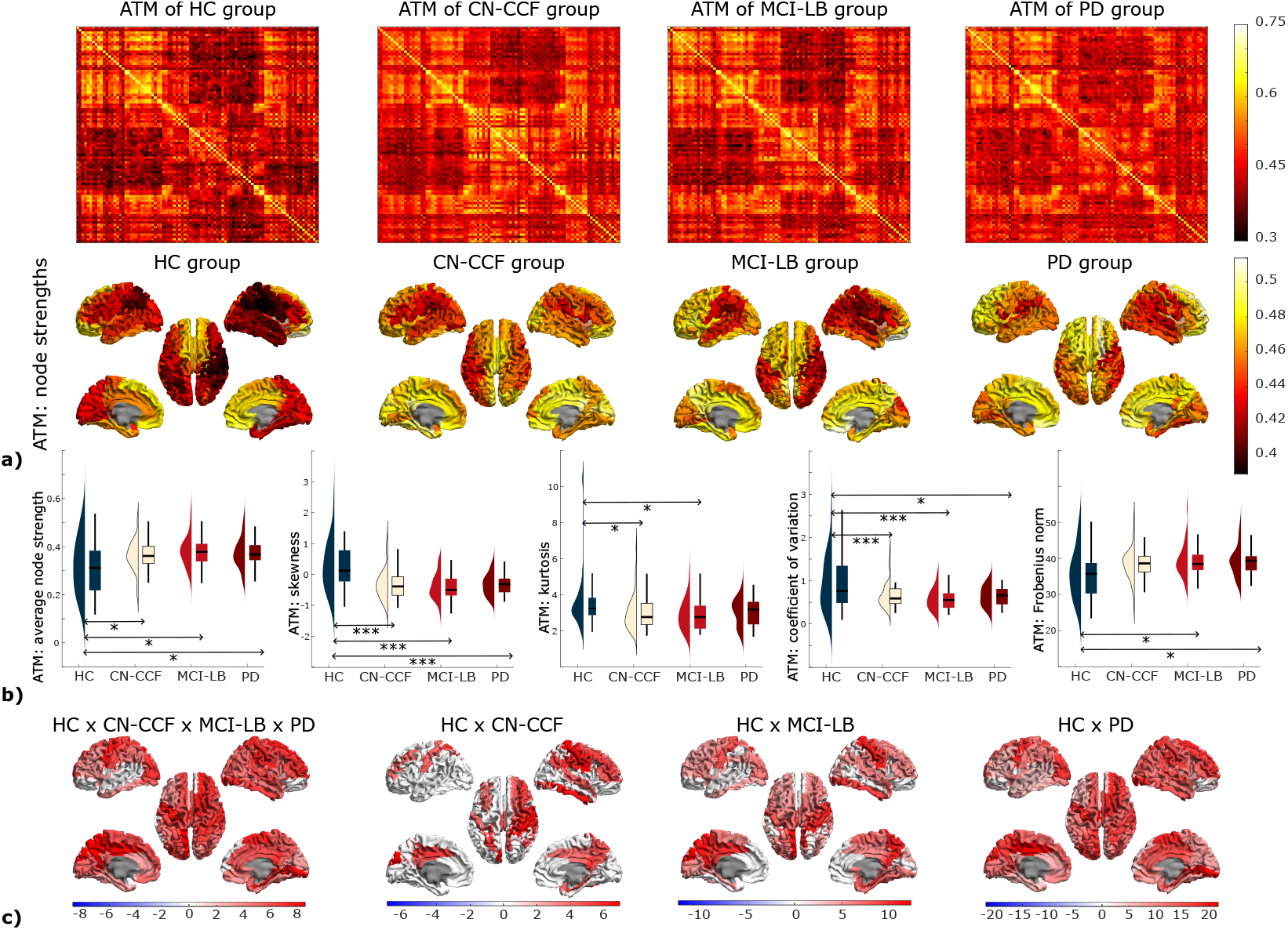
ATMs characteristics. **a)** Group-specific ATMs visualized as matrices (1^st^ row) and in brain space as node strengths (2^nd^ row). In the matrix form, regions are sorted from the top and left to the bottom and right as Frontal (1–30), Cingulate (31–36), Mesiotemporal (37–42), Occipital (43–56), Parietal (57–70), Subcortical (71–78), and Temporal (79–90), with left and right regions in alternating order. **b)** Between-group comparisons of global ATM’s features: average node strength, skewness, kurtosis, coefficient of variation, and Frobenius norm. * *p* < .05, uncorrected, *** *p* < .05, FDR corrected. **c)** F-statistics of between-group comparisons of ATM’s average node strengths. Only F-values with *p* < .05 are color-coded; note: red = average node strength is higher in the patient groups compared to HC.

**Fig. 4.**
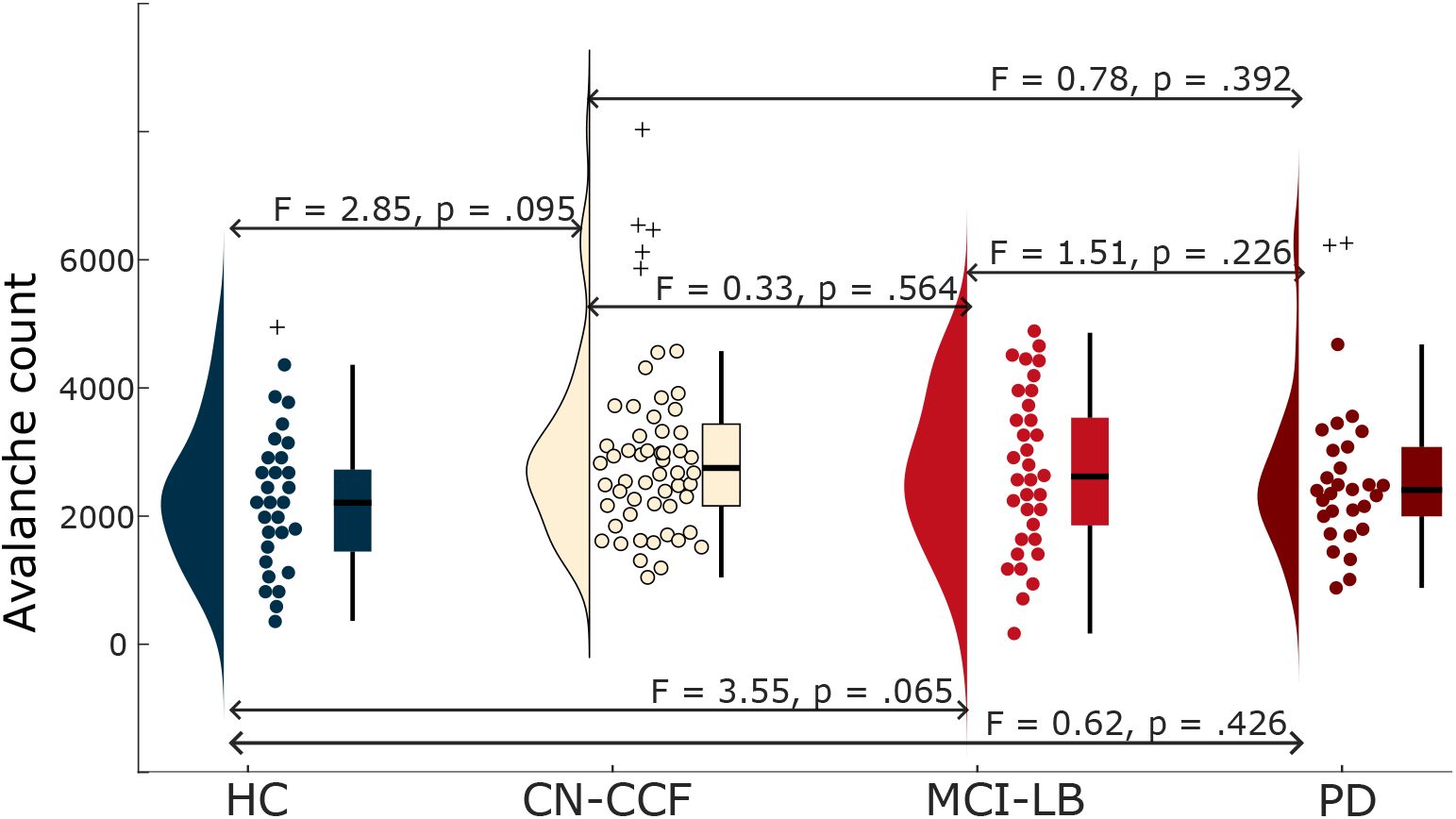
Group differences in avalanche count. Note: Crosses (+) mark 8 outliers that were removed before statistical analysis; n(HC) = 29, n(CN-CCF) = 51, n(MCI-LB) = 34, n(PD)=28) without outliers. Effects of age, sex, and LEDD were regressed out before analyses.

**Fig. 5.**
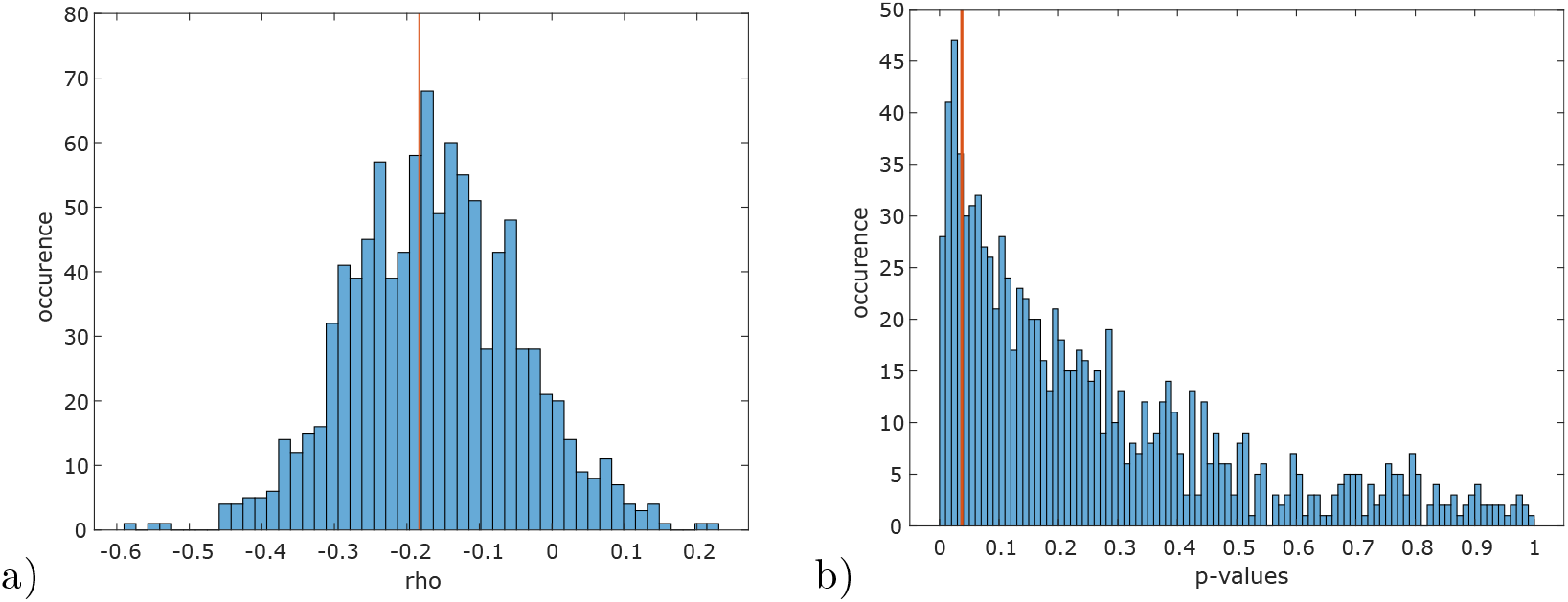
Relationship between flexibility and the rcLC. Bootstrap Spearman correlations computed 1,000-times on 50 randomly selected subjects, age and sex regressed out. A histogram of bootstrap correlations’ a) rhos and b) p-values. Orange line: the significance of the full result (rho = −.2 (CI[−.39, −.00]), *p* = .036 (CI [.005,.885])).

**Fig. 6.**
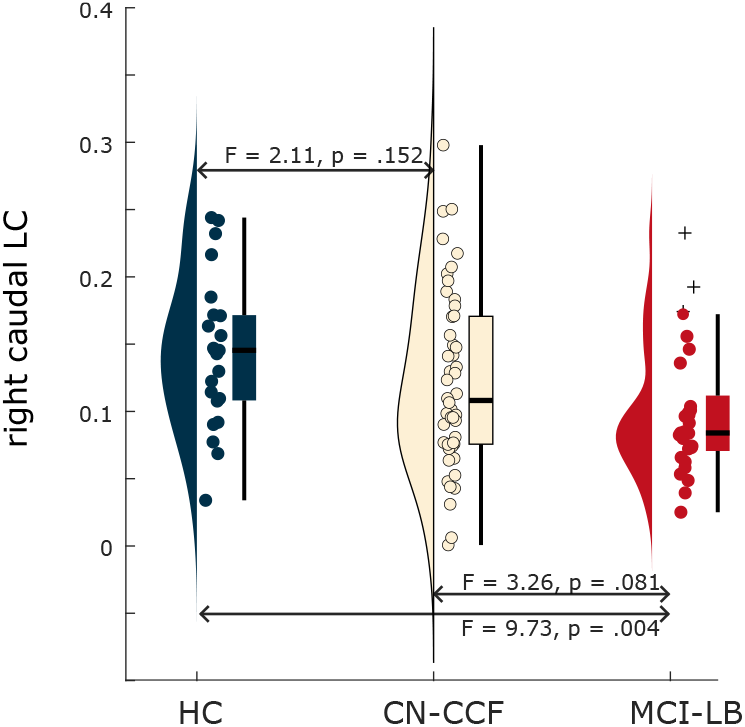
Group differences in the rcLC. n(HC) = 23, n(CN-CCF) = 48, n(MCI-LB) = 29. CI(HC x CN-CCF) [0.00, 5.34], CI(HC x MCI-LB) [0.00, 5.39], CI(CN-CCF x MCI-LB) [0.00, 5.55].

Individual ATMs were quantified using five global features: average node strength, skewness, kurtosis, coefficient of variation, and Frobenius norm; between-group differences are described in Table 3 and Fig 3b. On the regional level, node strengths showed significant differences between groups; their spatial patterns are visualized in Fig 3c.

**Table 3.**
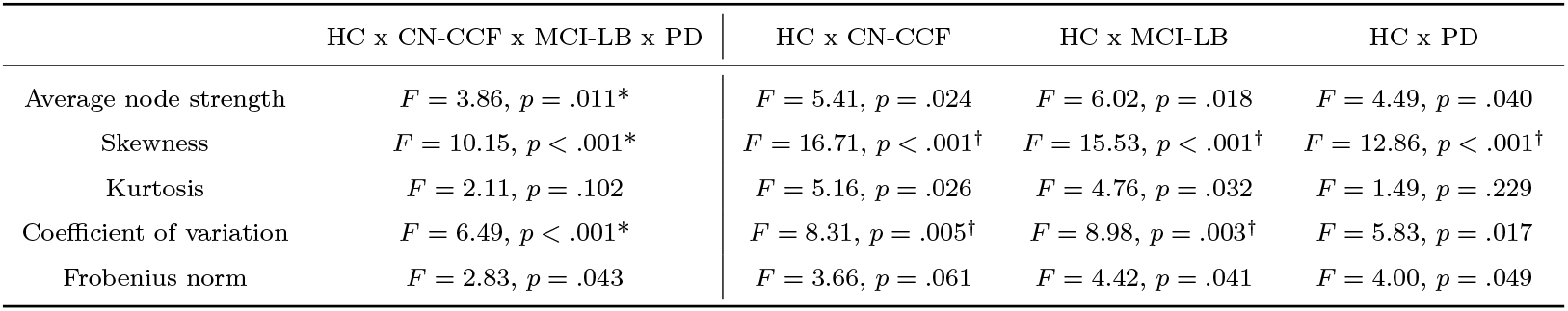
Between-group differences in ATMs characteristics, excluding outliers. * *p* < .05, FDR corrected within the omnibus tests comparing all four groups; ^†^ *p* < .05, FDR corrected within the post-hoc tests. Confidence intervals for all omnibus tests range between [.02, 3.92] and for post-hoc tests between [0, 5.48].

|  | HC x CN-CCF x MCI-LB x PD | HC x CN-CCF | HC x MCI-LB | HC x PD |
| --- | --- | --- | --- | --- |
| Average node strength | $F = 3.86, p = .011^*$ | $F = 5.41, p = .024$ | $F = 6.02, p = .018$ | $F = 4.49, p = .040$ |
| Skewness | $F = 10.15, p < .001^*$ | $F = 16.71, p < .001^\dagger$ | $F = 15.53, p < .001^\dagger$ | $F = 12.86, p < .001^\dagger$ |
| Kurtosis | $F = 2.11, p = .102$ | $F = 5.16, p = .026$ | $F = 4.76, p = .032$ | $F = 1.49, p = .229$ |
| Coefficient of variation | $F = 6.49, p < .001^*$ | $F = 8.31, p = .005^\dagger$ | $F = 8.98, p = .003^\dagger$ | $F = 5.83, p = .017$ |
| Frobenius norm | $F = 2.83, p = .043$ | $F = 3.66, p = .061$ | $F = 4.42, p = .041$ | $F = 4.00, p = .049$ |

**Table 4.**
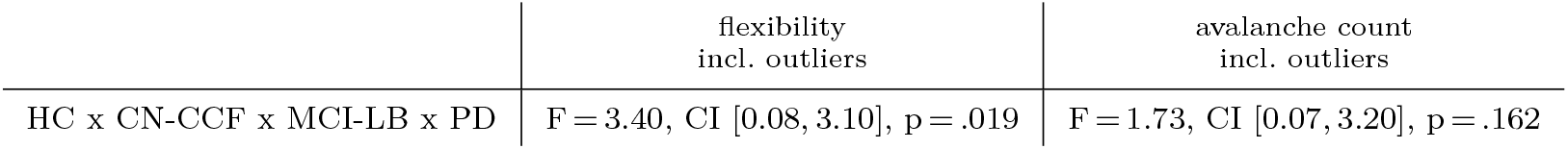
Between-group differences in flexibility and avalanche count, outliers included. Effects of age, sex, and LEDD were regressed out before analyses.

**Table 5.**
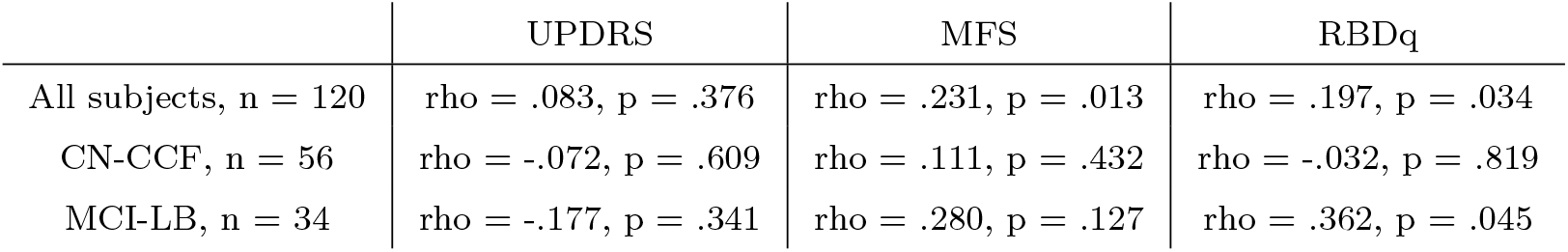
Partial Spearman’s correlations between flexibility and the core clinical features of interest. Age, sex, and depressive symptoms (Geriatric Depression Scale) as covariates.

Spearman’s correlations (age, sex, and LEDD regressed out) were conducted to examine associations between ATM’s features and flexibility. Of the five ATM features, skewness and coefficient of variation (CV) were significantly and negatively related to flexibility (*p* < .05, FDR corrected; n = 150; rho = −.45, *p* < .001, rho = −.22, *p* = .007, respectively). Given that ATMs were more negatively skewed in patient groups than in HC (rightward shift in the histograms pointing to greater prevalence of higher values), these results suggest that higher flexibility is associated with higher avalanche transition probabilities and lower variability (captured by CV).

## 3 Discussion

Our work investigated flexibility of large-scale neuronal avalanches in LBD, a subset of the neural *α*-synuclein diseases, as compared with healthy older adults. We found higher flexibility in two prodromal DLB groups (MCI-LB and CN-CCF) relative to HC, indicating that prodromal DLB patients exhibit a broader repertoire of distinct avalanche patterns than HC, while they did not differ in overall count of supra-threshold bursts of activity. This contrasts findings in several neurodegenerative conditions where flexibility was observed to be reduced [10–12, 32]. One plausible interpretation is that increased flexibility reflects richer repertoire of activity expressed across the EEG recording session, capturing dynamic fluctuations in cognition, including lapses of attention and arousal [33]. Alternatively, early DLB produces network dysregulation [34] or altered neuromodulatory tone (for example via rcLC degeneration), which transiently increases spontaneous network reconfiguration before the repertoire collapses as the disease progresses into DLB or PD dementia. This is in line with previous reports that temporal variability (that is, fluidity) sometimes reflects compensatory dynamics rather than disease severity [19]. While both explanations are plausible, we cannot draw further conclusions from our data, as our cohort did not include a group with fully developed DLB.

Furthermore, we observed that PD patients had higher flexibility than HC, however less so than the prodromal DLB groups, despite the fact that PD patients were advanced in the disease progression with relatively high LEDD, symptoms of cognitive impairment, symptoms of RBD, and cognitive fluctuations. This contrasts the study by Sorrentino et al., who observed a significantly reduced repertoire of avalanches in PD patients compared to HC [11]. This difference may be attributable to two factors. First, the present study acquired data in the medication-ON state, whereas Sorrentino et al. [11] assessed flexibility during the OFF state. Second, our cohort exhibited higher mean LEDD (840 mg vs. 289.9 mg in [11]), and already manifested symptoms of REM sleep behavior disorder and cognitive fluctuations, indicating more advanced PD progression [6].

Taken together, increased flexibility in LBD cohort may indicate the presence of malignant non-motor symptoms of LBD, which are associated with higher risk of phenoconversion to DLB or PD dementia [6]. This interpretation also aligns with our observation that asymmetry in the distribution or spatial spread of flexibility was not associated with motor impairment in PD patients, even though *α*-synuclein pathology in PD is frequently asymmetric [35, 36]. An alternative, but complementary, interpretation is consistent with the *Synuclein Origin and Connectome model* (SOC model [37]). Specifically, flexibility may indicate a body-first progression of *α*-synuclein, as it was positively associated with fluctuations in cognition and symptoms of REM sleep behavior disorder, but not with parkinsonism. Moreover, it was associated with structural integrity of the caudal part of LC but not with the structural integrity of SNpc in prodromal DLB groups. In advanced fully developed PD patients, the flexibility was not associated with PD symptoms or with flexibility’s asymmetry while it was associated with their malignant non-motor symptoms predicting progression to dementia and suggesting a spread of *α*-synuclein already both to association cortex and to caudal parts of the brain stem.

Further, we hypothesized that flexibility, reflecting the richness of brain dynamics, would be associated with the structural integrity of neuromodulatory nuclei. LC, a small noradrenergic nucleus in the pons, provides widespread cortical noradrenergic modulation and is therefore an essential driver of several attentional and executive functions, including orienting attention, vigilance, stress responses [38], arousal, or sustained attention [39]. LC activity strongly influences the dynamics of brain states in healthy individuals [40], and altered temporal variability in functional connectivity (i.e., fluidity) in prodromal DLB has been linked to decreased integrity of the rcLC [19]. In the present study, increased flexibility was associated with decreased integrity of the rcLC, but not with SNpc integrity. This finding further supports the relevance of the rcLC in the body-first pathology, as multiple studies have reported total or sub-regional LC involvement in this subtype. For instance, Passaretti et al. [35] reported decreased caudal LC-cortical connectivity in the body-first patients, whereas rostral LC-cortical connectivity was more affected in the brain-first *α*-synuclein aggregation. Others reported overall decreased LC signal intensity [36] and decreased LC-pons ratio [41] in the body-first subtype. The SNpc, which produces dopamine and projects to the striatum to regulate movement and learned responses, is primarily affected in the brain-first subtype [42, 43].

Analyses of spatial avalanche patterns (ATM) revealed higher transition probabilities in prodromal LBD compared to HC, suggesting breakdown of brain’s dynamic communication. Avalanches propagated more readily specifically through the medial prefrontal, orbitofrontal, and temporal cortices, extending into inferior frontal and posterior parietal areas - regions implicated in cognitive fluctuations, executive dysfunction, and visuospatial deficits characteristic of prodromal DLB [44].

Although our study uses a multimodal approach to investigate LBD, several limitations should be acknowledged. First, despite employing robust statistical methods, the relatively small group sizes may limit the generalizability of our findings. Additionally, PD participants were recruited under a separate protocol, resulting in the MRI data (LC and SNpc imaging) that were not comparable between PD and other subgroups. The LC and SNpc were imaged using different MRI sequences - neuromelanin-sensitive and free-water imaging, respectively. This choice reflects the fact that the LC is too small for reliable free-water imaging, and neuromelanin-sensitive sequences for the SNpc were not yet available at our core facility. Another limitation is the absence of *α*-synuclein–specific biomarkers, such as seed amplification assays, which are not yet available in our region. The lack of these pathological markers limits diagnostic certainty and precludes definitive classification of participants according to underlying *α*-synuclein pathology. Finally, although the RBD questionnaire provides useful screening information, it does not replace objective polysomnographic assessment. Replication of our findings using standardized diagnostic procedures is therefore warranted.

In conclusion, this multimodal study combined high-density EEG with MRI imaging of LC and SNpc to examine how spontaneous cascades of neuronal activity relate to the integrity of catecholamine-rich nuclei and clinical symptoms. We observed greater flexibility in subjects with LBD, which was positively associated with cognitive fluctuations and symptoms of REM-sleep behavior disorder, but not with the severity of parkinsonism, suggesting a phenotype-specific alteration linked to the fluctuations in alertness. This interpretation is further supported by the inverse relationship with the rcLC integrity, indicating that dysregulated dynamics may arise from early LC dysfunction rather than SN pathology. Finally, by associating large-scale brain dynamics and neuromodulatory-nuclei integrity across the LBD phenotypes, we propose that flexibility may reflect *α*-synuclein spread which, if confirmed by *α*-synuclein detection, aligns with the SOC model [37].

## 4 Methods

### 4.1 Sample

Participants were recruited through advertisements and screened via telephone for inclusion and exclusion criteria. Inclusion criteria included age above 50 years, subjective symptoms of prodromal DLB [1], or PD diagnosis (criteria by Postuma et al. [45]). Exclusion criteria included severe medical conditions, major brain injury, other psychiatric or neurological disorders, dementia, alcohol or drug abuse, and MRI contraindications. Three subjects were excluded due to incomplete neuropsychological assessment, nine subjects due to clinical diagnoses (e.g., progressive supranuclear palsy, or post-stroke apraxia), and fifteen with MCI were excluded because they did not manifest the core clinical features of prodromal DLB.

After the phone interview, neuropsychologists and a neurologist conducted clinical and neuropsychological evaluations with HC and prodromal LBD subjects, based on which participants were classified as HC or prodromal LBD, PD were diagnosed by neurologists. Subsequently, subjects underwent high-density resting-state EEG, DTI, and neuromelanin-sensitive MRI across two visits. PD subjects did not undergo DTI nor neuromelanin-sensitive MRI).

The final sample (n = 150) comprised of HC = 30, PD = 30, and prodromal LBD subjects with (MCI-LB; n = 34) or without (CN-CCF; n = 56) cognitive impairment. The following criteria were used to classify the subjects:

#### Noncognitive symptoms

Diseases included in LBD spectrum are diagnosed based on the sequence, severity, and predominance of clinical signs and symptoms across disease progression [46]. Subjects were classified as prodromal LBD if they had at least one of the core clinical features of prodromal DLB, as defined by McKeith et al. [1], none of the subjects had any clinical diagnosis. These features were operationalized using the following scales and thresholds: Unified Parkinson’s Disease Rating Scale, part 3 [47] (UPDRS; cut-off ≥ 4 points); Neuropsychiatric Inventory Questionnaire-hallucination subscale [27] (NPI; cut-off = 1 point); Mayo Fluctuations Scale [25] (MFS; cut-off ≥ 3 points); REM Sleep Behavior Disorder Questionnaire [26] (RBDq; cut-off ≥ 5 points).

#### Cognitive symptoms

subjects were classified as MCI-LB if their performance on two or more cognitive tests was below −1 SD of the age-appropriate norm (see Mitterová et al. [19] for details on neuropsychological battery).

This study was approved by the Masaryk University Research Ethics Committee (approval numbers EKV-2022-094 and EKV-2021-069) and conducted in accordance with the Declaration of Helsinki and relevant ethical guidelines. All participants provided written informed consent before data acquisition. Clinical trial number: not applicable.

### 4.2 EEG processing

#### 4.2.1 EEG preprocessing

Fifteen minutes of resting state, eyes closed, were recorded using high density scalp EEG system (HD-EEG Electrical Geodesics, Inc.; EGI GES 400 MR) with 256 channels, 1 kHz sampling frequency, and reference electrode set to Cz channel. The processing was done using MATLAB, Fieldtrip [48], EEGLAB [49], and Cartool [50] similarly as in [19, 51]. Data were reduced to 204 electrodes by discarding facial and neck electrodes [52], filtered to 0.1-100 Hz, implementing a second-order Butterworth filter, 12 dB/octave roll-off, and forward and backward passes [53], and downsampled four times to the sampling frequency Fs = 250 Hz. Bad channels were automatically detected and removed (a channel was considered as bad if its maximum amplitude was below 10 or exceeded the mean maximum amplitude across all channels by more than three standard deviations). Afterwards, artifact segments were automatically detected if their global field power (gfp) exceeded ten standard deviations above average of all gfps). The ICA decomposition (RUNICA) was run and artifact components were manually detected (eye movement, eye blinks, cardio; max 3-5 components) [54]. After the removal of these components and automatic interpolation of previously detected bad channels, accompanied by a manual check, the first 7 minutes of signals were rereferenced to average and source reconstructed [55, 56] into 116 regions (ROIs) of AAL atlas [57] using a template transformation matrix. Cerebellar and vermis ROIs were discarded and 90 ROIs kept. After the last manual detection of artifact segments, the signals were filtered into the broadband frequency band of interest (0.1–100 Hz, the lower cut-off kept the same as in other EEG studies evaluating neuronal avalanches [15, 58, 59]), and the artifact segments were cut out.

#### 4.2.2 Avalanche detection

The processing was done similarly as in [11, 14, 16]. Data lengths were computed across subjects, and all signals were limited to the same length (time cut = 96,471 timepoints ≅ 6.43 min). The distribution of the z-scored signal was inspected by computing and visualizing its histogram (Supplementary Fig 7a). A detection threshold of 3 standard deviations was selected based on the point at which the empirical distribution deviates from a Gaussian profile, indicating the onset of non-Gaussian, large-amplitude events. Neuronal avalanches were subsequently detected independently within each segment using this threshold. To verify that the detected events were consistent with the definition of neuronal avalanches—namely, that they form cascades whose size and duration distributions exhibit scale-free behavior following a power-law [9]—the distributions of avalanche size and duration were computed after concatenating events across subjects. Both distributions displayed power-law–like behavior as shown in Supplementary Fig 7b,c. Therefore, these events can be referred to as neuronal avalanches. The consequent computations of flexibility, see below, were repeated on different thresholds to show the stability of findings presented here; see Supplementary Table 6.

**Table 6.** The influence of threshold selection on avalanche detection. Each row shows a given threshold, median (25^th^, 75^th^ quartiles) of flexibility across subjects (*n* = 119, excluding the outlier visible in Fig 7), and Spearman’s correlations. The correlations yielded strong relationships between flexibility computed on different thresholds, and support the stability of between-group differences in flexibility over thresholds. The threshold *thr* = 3 (in dark blue) is the one chosen for the analyses in this manuscript.

| | Median (Q1, Q3) | $thr = 3$ | $thr = 2.5$ | $thr = 2$ |
| --- | --- | --- | --- | --- |
| $thr = 3.5$ | 314 (220, 413) | $\rho = .86, p < 10^{-36}$ | $\rho = .68, p < 10^{-17}$ | $\rho = .71, p < 10^{-20}$ |
| $thr = 3$ | 693 (514, 944) | — | $\rho = .94, p < 10^{-55}$ | $\rho = .92, p < 10^{-48}$ |
| $thr = 2.5$ | 1670 (1149, 2460) | — | — | $\rho = .95, p < 10^{-59}$ |
| $thr = 2$ | 2843 (1919, 3901) | — | — | — |

**Fig. 7.**
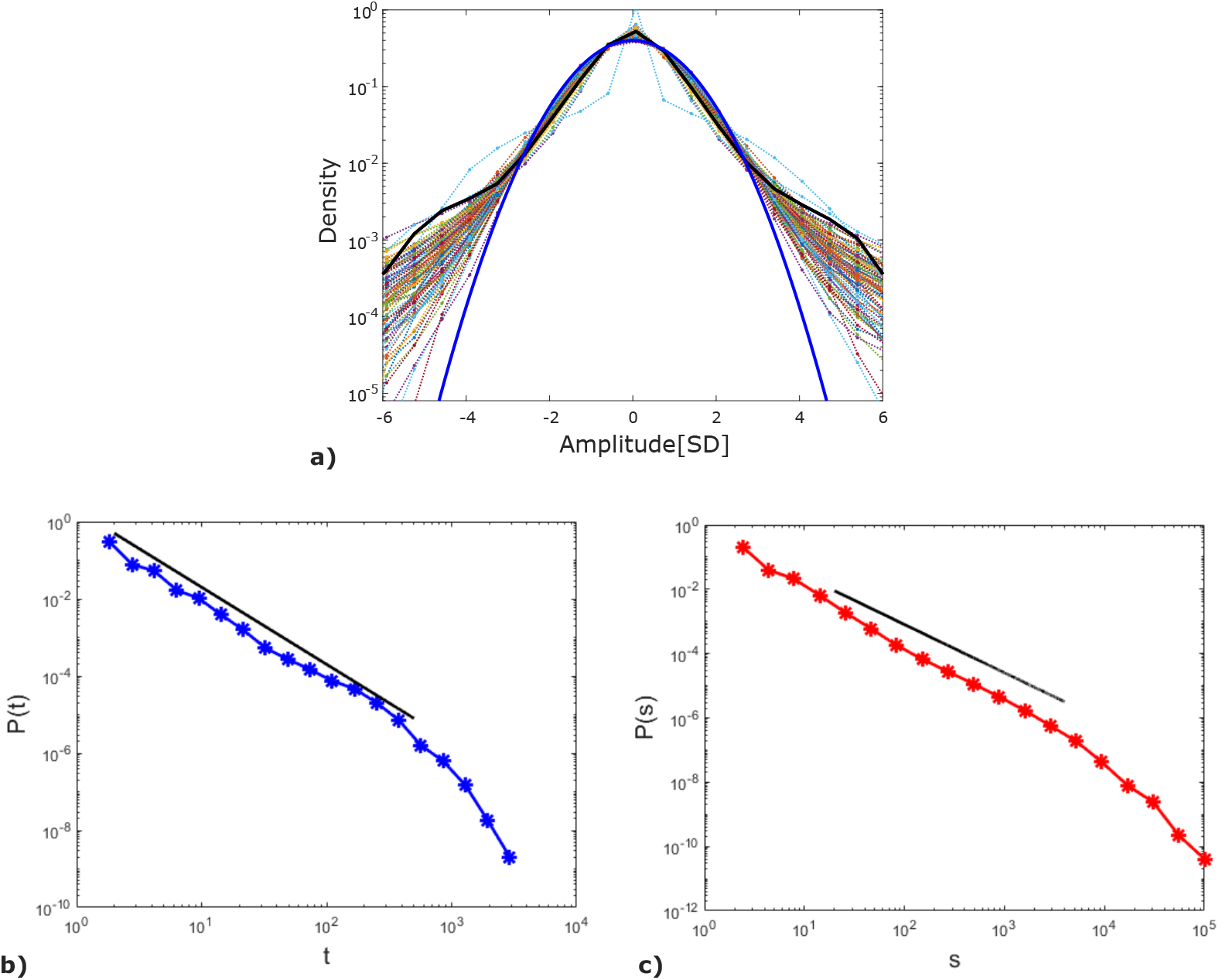
Signal distribution for avalanche threshold detection. *n* = 150, **a)** semilog plot of signals distributions for each subject. Based on this plot, *thr* = 3 was chosen as a threshold for avalanche detection (real mean signals distributions visibly cross the simulated normal distribution (in blue)). One outlier—a healthy control (visually distinguished in teal in the plot)—was identified. As PERMANOVA inherently accommodates outliers, this subject was included in all analyses reported in the manuscript. For completeness, all analyses were also repeated after excluding this individual, and the results remained consistent; **b)** supra-threshold events duration; **c)** size distributions. The straight lines represent the slopes expected from the theoretical exponents of the branching process.

#### 4.2.3 Quantifying EEG dynamics

EEG dynamics was quantified by the total number of avalanches, flexibility, flexibility’s hemispheric symmetry, avalanche transition matrix and features derived from it. Here, these variables are explained.

**Total number of avalanches** is the count of active (over the threshold) sequences. A sequence (= an avalanche) begins when at least one channels becomes active and ends when all channels become inactive in the following consecutive time bins. An avalanche is represented by its spatial pattern informing about ROIs involved in the avalanche.

**Flexibility** was computed as a number of unique binarized avalanche patterns per subject.

##### Flexibility’s hemispheric symmetry

Considering only avalanches propagating unilaterally (cross-hemispheric avalanches were discarded), flexibility was computed separately for the left and right hemisphere. A ratio quantifying hemispheric symmetry in flexibility was computed as *flexiratio* = *abs*((*flexi*_*L*_ − *flexi*_*R*_)*/*(*flexi*_*L*_ + *flexi*_*R*_)). Here, *flexiratio* = 0 indicates complete symmetry, while *flexiratio* = 1 indicates full asymmetry. Symmetry implies that both hemispheres exhibit functional repertoires of similar size in terms of variability in avalanche patterns.

**Avalanche transition matrix (ATM)** is an *N × N* matrix, where *N* is the number of ROIs, and *ATM*_*i,j*_ is a probability that channel *j* was active at time *t* + 1 while channel *i* was active at time *t*. That is, the ATM represents probabilities that channels *i* and *j* were involved in the same avalanches.

In the present work, following [17], we quantified the ATMs using five features: average node strength, defined as 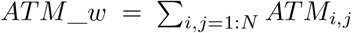 skewness, which captures the asymmetry of the ATM distribution (positive values indicate a leftward shift, negative values a rightward shift); kurtosis, which characterizes the tailedness of the ATM distribution (positive values indicate a high peak with light tails, whereas negative values indicate a flatter peak with heavier tails); the coefficient of variation defined as 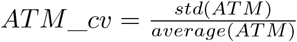 and Frobenius norm defined as 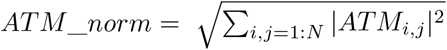, which quantifies the scalar size of the ATM.

### 4.3 MRI processing

Imaging was performed on a 3 T Siemens Prisma scanner at CEITEC, Masaryk University, Brno, Czech Republic. T1 imaging was performed with MPRAGE sequence with 240 sagittal slices, TR = 2400 ms, TE = 2.27 ms, FOV = 218 mm, FA = 8°, matrix size 192 x 256 x 256, isotropic voxel 0.85 mm. The MRI protocol included a neuromelanin-sensitive sequence targeting the LC and free-water imaging assessing microstructural integrity of the SNpc. Free-water imaging could not be applied to the LC due to insufficient spatial resolution of the diffusion-weighted imaging sequence.

#### 4.3.1 Neuromelanin-sensitive MRI acquisition: Locus coeruleus processing

LC-focused neuromelanin-sensitive MRI (NMS-MRI) data was processed following the same pipeline used in the prior work [19, 21, 60]. T1-weighted images were first skull-stripped using ANTs [61] and spatially normalized to MNI space. Each NMS-MRI volume was then rigidly co-registered to the native-space T1-weighted image, and all transformations were concatenated to warp the LC and pontine tegmentum (PT) masks in a single step.

LC signal extraction adhered to established procedures. An overinclusive, manually-defined mask specified the LC search region, while the reference region was placed in the central PT due to its highly stable signal-to-noise characteristics. This enabled normalization of intensity across participants and slices. Intensities were scaled such that the PT reference signal was equivalent across all images, and the LC contrast ratio (*LC_CR*) was computed as *LC_CR = (LC intensity − PT intensity) / PT intensity*.

For each hemisphere, the voxel with the maximal intensity within the LC search area was identified, and a five-voxel cross-shaped kernel was centered on this peak to capture local signal. When the peak voxel was adjacent to the fourth ventricle, the kernel was shifted posteriorly by one voxel to avoid partial-volume contamination. Signals were extracted from three axial slices spanning the rostro-caudal extent of the LC, with the middle slice positioned 7 mm inferior to the lower border of the inferior colliculus, following prior anatomical work.

In the present study, the analysis focused on the right caudal LC, consistent with previous findings that this region shows selective signal reduction in prodromal DLB relative to healthy controls and that this reduction relates to cognitive performance [19, 21].

#### 4.3.2 Free-water DTI imaging: Substantia nigra processing

Interleaved multi-slice DWI was performed using the following parameters: b-values 0, 500, 1000, 1750, and 2000 s/mm^2^, with overall 133 non-colinear directions; TR = 3374 ms, TE = 73.6 ms, FOV = 210 mm, acquisition matrix 140 × 140 × 90, isotropic voxel 1.5 mm. The same sequence was acquired with opposite phase polarity.

Diffusion MRI data were preprocessed using MRtrix 3.0 [62] and FSL 6.0.1 [63]. In short, data were corrected for noise, Gibbs ringing artifacts, motion, eddy current, and susceptibility distortions. Also bias field correction was performed in Advanced Normalisation Tool (ANTs; [61]). After preprocessing, volumes with b-values 0, 500 and 1000 s/mm^2^ were extracted and underwent the free-water elimination (FWE-DTI) model in DIPY [64, 65]. The metric of interest was the volume fraction of free-water (FW).

The FWE-DTI model is a two-compartment model, therefore is well posed for multi-shell data. We used the lower b-values because the model is less affected by non-Gaussian diffusion effects [66]. Mask of SNpc was obtained from Zhang et al. [67], by merging the masks of lateral and medial SN. Registration of the masks from MNI to individual T1 and DWI space was performed using nonlinear registration in ANTs.

### 4.4 Statistical analyses

#### 4.4.1 Between-group differences

For testing between-group differences of cross-correlated non-gaussian variables that are derived from the brain data, the permutational multivariate analysis of variance (PERMANOVA) conducted in MATLAB [29] was adopted, with age and sex regressed out of all variables beforehand. The distance matrix, serving as an input into the PERMANOVA, was computed using Mahalanobis distance due to its ability to deal with correlated data. The significance of between-group differences was assessed using PERMANOVA, with F-statistics and permutation-based p-values derived from 20,000 permutations.

#### 4.4.2 Associations with clinical features

Spearman’s correlations were conducted to examine relationships between EEG features (flexibility and flexibility asymmetry index) and neuromodulatory nuclei (rcLC and SNpc) and core clinical features in the combined sample and separate subgroups. Core clinical features of prodromal DLB, following McKeith et al. [1] were operationalized as the Mayo Fluctuation scale (MFS, A shortened 4-item version of the MFS asks informants to report on cognitive fluctuations in daily life), the REM Sleep Behavior Questionnaire (RBDq, assessing symptoms of REM sleep behavior disorder), and the Unified Parkinson’s Disease Rating Scale (UPDRS, assessing symptoms of parkinsonism); The Neuropsychiatric Inventory for visual hallucinations was not included in the correlations as it consists of only two items. The correlations included the following covariates: age, sex, LEDD, and depressive symptoms (Geriatric Depression Scale score), as indicated in the results. Additionally, ATM features were correlated with flexibility to relate the breadth of the functional repertoire to the global propagation of avalanches across the whole brain.

## 5 Acknowledgements

We acknowledge the contribution of the core facility MAFIL supported by MEYS CR (LM2023050 Czech-BioImaging), part of the Euro-BioImaging (www.eurobioimaging.eu) ALM and Medical Imaging Node (Brno, CZ). We thank Anne Johnson for the English editing and Vanessa Skyblová for her help with participants.

## Declarations

### Data and code availability

Data and code are available upon reasonable request.

### Author contribution

**E.V.**: Writing – original draft, Writing – review & editing, Visualization, Methodology, Investigation, Formal analysis, Data curation, Conceptualization. **K.M**.: Writing – original draft, Writing – review & editing, Visualization, Methodology, Investigation, Formal analysis, Data curation, Conceptualization. **M.A**.: Writing – review & editing, Methodology, Data curation. **A.K**.: Writing – review & editing, Data curation. **A.Š.M**.: Writing – review & editing, Data curation. **L.B**.: Writing – review & editing, Data curation. **M.L**.: Writing – review & editing, Data curation. **I.R**.: Writing – review & editing, Supervision, Funding acquisition. **P.S**.: Writing – review & editing, Supervision, Methodology. **J.F**.: Writing – review & editing, Supervision, Methodology, Conceptualization, Funding acquisition.

## 6 Supplementary information

